# The Human Microglia Atlas (HuMicA) Unravels Changes in Homeostatic and Disease-Associated Microglia Subsets across Neurodegenerative Conditions

**DOI:** 10.1101/2023.08.01.550767

**Authors:** Ricardo Martins-Ferreira, Josep Calafell-Segura, Bárbara Leal, Javier Rodríguez-Ubreva, Elisabetta Mereu, Paulo Pinho e Costa, Esteban Ballestar

## Abstract

Dysregulated microglia activation, leading to neuroinflammation, is crucial in neurodegenerative disease development and progression. The initial M1/M2 dual activation classification for microglia is outdated. Even the ‘disease-associated microglia’ (DAM) phenotype, firstly described in mice, falls short in representing the diverse microglia phenotypes in pathology. In this study, we have constructed a transcriptomic atlas of human brain immune cells by integrating single-nucleus (sn)RNA-seq datasets from multiple neurodegenerative conditions. Sixteen datasets were included, comprising 295 samples from patients with Alzheimer’s disease, autism spectrum disorder, epilepsy, multiple sclerosis, Lewy body diseases, COVID-19, and healthy controls. The integrated *Human Microglia Atlas* (*HuMicA*) dataset included 60,557 nuclei and revealed 11 microglial subpopulations distributed across all pathological and healthy conditions. Among these, we identified four different homeostatic clusters as well as pathological phenotypes. These included two stages of early and late activation of the DAM phenotype and the disease-inflammatory macrophage (DIM) phenotype, which was recently described in mice, and is also present in human microglia, as indicated by our analysis. The high versatility of microglia is evident through changes in subset distribution across various pathologies, suggesting their contribution in shaping pathological phenotypes. Our analysis showed overall depletion of four substates of homeostatic microglia, and expansion of niche subpopulations within the DAM and DIM spectrum across distinct neurodegenerative pathologies. The *HuMicA* is invaluable in advancing the study of microglia biology in both healthy and disease settings.

## INTRODUCTION

Microglia are central immune mediators in the brain. They originate from myeloid precursors and migrate from the yolk sac to the brain parenchyma during the embryonic stage^1,2^. In the brain of healthy individuals, microglia orchestrate a wide network of processes essential for the development and maintenance of the normal functioning central nervous system (CNS), including modulation of neuronal connectivity through synaptic remodelling and structural organization, myelination, blood-brain barrier integrity and vasculogenesis. In pathological contexts, microglia generally acquire an activated state, transitioning from a resting elongated and ramified form to an amoeboid macrophage-like morphology, accompanied by an overall enhanced pro-inflammatory phenotype^3^. Despite being a normal physiological mechanism, unbalanced and exacerbated microglia activation has consistently been demonstrated to contribute to the development and progression of neurological diseases.

The landscape of microglia activation was initially described by the M1/M2 polarization, wherein the M1 pro-inflammatory phenotype was considered neurotoxic and the M2 tolerant phenotype was thought to be neuroprotective^4^. In 2017, Keren-Shaul *et al*. described what is still regarded as the consensus microglia pathological phenotype, known as the “disease-associated microglia” (DAM). This milestone was achieved through the application of single-cell (sc) transcriptomics in a mice model of Alzheimer’s disease (AD). The authors identified phenotypically distinct subpopulations of microglia that were enriched in pathological conditions. These subpopulations exhibited downregulation of homeostatic microglia markers and activation of pathways involved in the immune response and phagocytosis^5^.

The implementation of single-nucleus (sn)RNA-seq methods has enabled the analysis of frozen tissue samples at the cellular level^6^, thereby maximizing the utilization of numerous previously frozen brain samples. This advancement has paved the way for human brain tissue studies across various neurological diseases, such as AD^7–12^, multiple sclerosis (MS)^13,14^, autism spectrum disorder^15^ and Lewy Body Diseases (LBD)^16^. Individual attention has been devoted to microglia, revealing evidence of dysregulated gene expression in pathology and, in certain cases, an enrichment of DAM traits. These findings suggest an inter-disease prevalence of this microglial phenotype. However, it is understandable that most of these studies place greater emphasis on the most abundant and traditionally considered more relevant cell types, particularly neuron populations.

It is becoming increasingly evident that the M1/M2 activation and the DAM phenotype are inadequate to fully comprehend the complexity of microglia behavior in both health and disease. It can be inferred that the plasticity of microglial function in homeostasis and injury/disease may follow a multi-subpopulation pattern influenced by the type of disease, sex, age and even spatial localization^17,18^.

In the human brain, the comprehensive characterization of microglia subpopulations is significantly limited by the low yield of microglia within snRNA-seq datasets. To address this limitation, we performed an *in-silico* integration of multiple snRNA-seq datasets from human brain tissue. These datasets included samples from patients diagnosed with AD, MS, autism, LBD and epilepsy, as well as samples collected from neurologically healthy individuals. Recent snRNA-seq studies demonstrated a pro-inflammatory DAM-like phenotype in severe COVID-19 brain tissue^19,20^. Considering these findings and the high incidence of long-term cerebral symptomatology associated with SARS-COV2 infection^21–23^, we deemed it relevant to include snRNA-seq data from severe COVID-19 patients.

In this study, we have successfully generated a comprehensive *Human Microglia Atlas (HuMicA)* consisting of 60,557 nuclei, derived from 295 individual subjects across sixteen datasets and seven clinical settings. Our results indicate that *HuMicA* is a unique resource and toolset that will facilitate further transcriptomic studies in human microglia in both health and disease.

## RESULTS

### Integration of the Human Microglia Atlas (HuMicA)

A total of sixteen publicly available snRNA-seq datasets on human brain tissue samples were utilized for this study (Table 1). These datasets encompassed samples from patients diagnosed with AD, autism, epilepsy, MS, LBD and severe COVID-19, as well as samples from individuals without any neurodegenerative diagnosis. Detailed information of each individual subject included in this study is available in Supplementary Table 1. The integration pipeline (Figure 1A) started by separately processing each dataset and annotating the main CNS cell types (neurons, oligodendrocytes, astrocytes, oligodendrocyte progenitor cells (OPCs), endothelial cells and immune cells) (Supplementary Figure 1 and Supplementary Figure 2) (see Methods). Subsequently, the immune cell clusters from each dataset were integrated, resulting in a final object comprising a total of 64,438 nuclei. We observed a uniform distribution of nuclei per cluster for all the studied variables, including brain region (Supplementary Figure 3A-F), despite the heterogeneity of the data used in relation to the source study, brain tissue region, age, pathological condition, gender and state of the tissue (post-mortem or surgical resection). This finding is consistent with previous data that suggests that microglia exhibit consistent patterns across different brain sections, unlike other CNS cell types^24^. We also assessed potential confounding effects and variance explained by each covariable. We observed that inter-subject variability is the main contributor to variance, while the other variables demonstrated a residual contribution (Supplementary Figure 3G).

**Figure 1.**
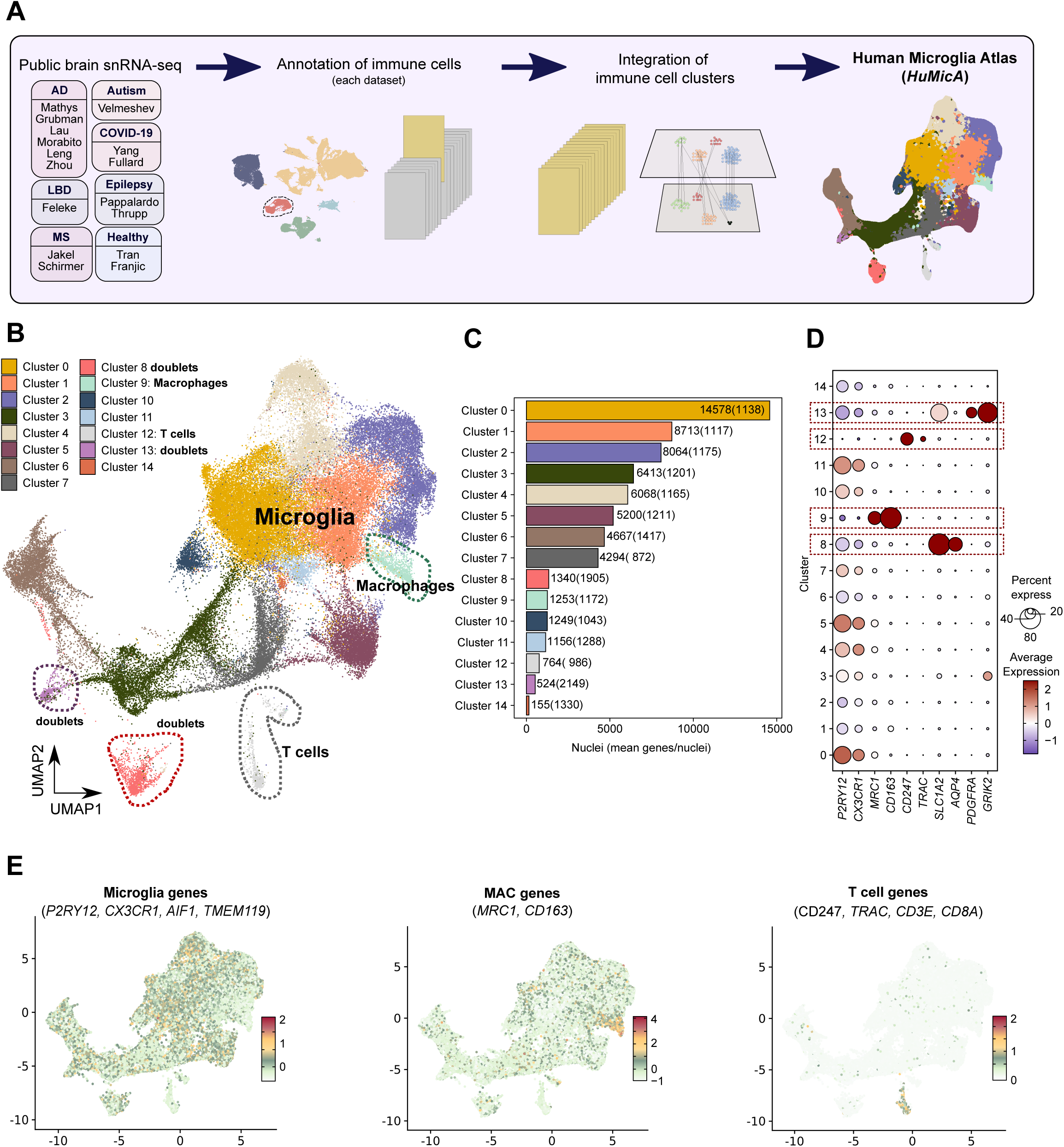
(A) Schematic overview of the pipeline used to obtain the integrated *HuMicA* (*Human Microglia Atlas*) object. **(B)** UMAP visualization of the integrated and clustered Seurat object, annotated by the fifteen obtained clusters. **(C)** Barplot representation of the number of nuclei and the mean genes detected per nuclei in each cluster. **(D)** Dot plot representation of the expression of canonical markers of microglia (*P2RY12*, *CX3CR1*), macrophages (*MRC1*, *CD163*), T cells (*CD247*, *TRAC*), astrocytes (*SLC1A2, AQP4*), OPCs (*PDGFRA*) and neurons (*GRIK2*) across all clusters. **(E)** UMAPs showing the module score (MSc) expression of microglia (*P2RY12*, *CX3CR1, AIF1, TMEM119*), macrophage (*MRC1*, *CD163*) and T cell (*CD247*, *TRAC, CDE3, CD8A*) markers.

**Table 1.**
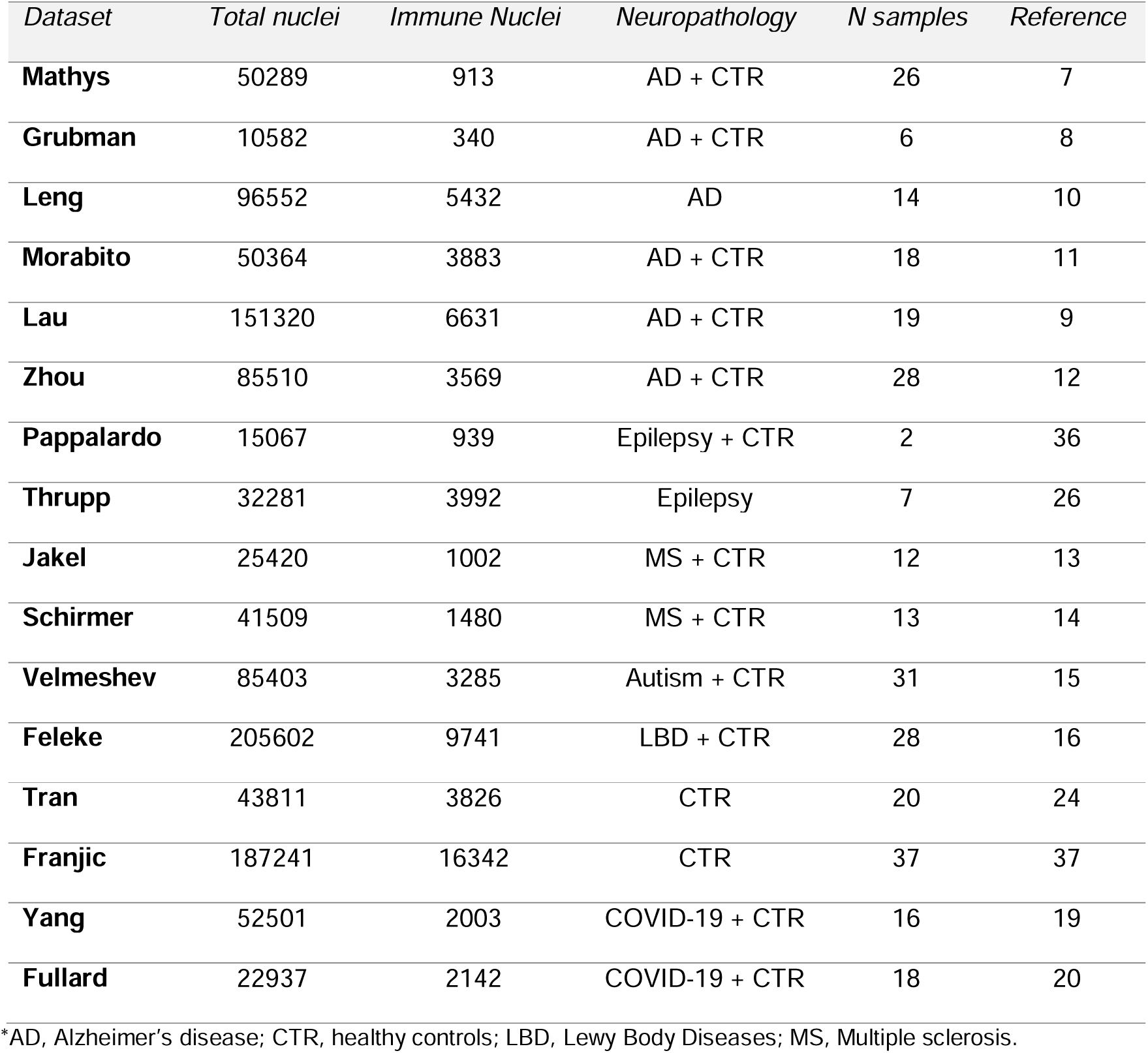
Summary of the public datasets used in this study, including the total number of nuclei after pre-processing, the number of immune nuclei, the neuropathological settings encompassed by each dataset and the number of individuals samples.

| *Dataset* | *Total nuclei* | *Immune Nuclei* | *Neuropathology* | *N samples* | *Reference* |
| --- | --- | --- | --- | --- | --- |
| **Mathys** | 50289 | 913 | AD + CTR | 26 | 7 |
| **Grubman** | 10582 | 340 | AD + CTR | 6 | 8 |
| **Leng** | 96552 | 5432 | AD | 14 | 10 |
| **Morabito** | 50364 | 3883 | AD + CTR | 18 | 11 |
| **Lau** | 151320 | 6631 | AD + CTR | 19 | 9 |
| **Zhou** | 85510 | 3569 | AD + CTR | 28 | 12 |
| **Pappalardo** | 15067 | 939 | Epilepsy + CTR | 2 | 36 |
| **Thrupp** | 32281 | 3992 | Epilepsy | 7 | 26 |
| **Jakel** | 25420 | 1002 | MS + CTR | 12 | 13 |
| **Schirmer** | 41509 | 1480 | MS + CTR | 13 | 14 |
| **Velmeshev** | 85403 | 3285 | Autism + CTR | 31 | 15 |
| **Feleke** | 205602 | 9741 | LBD + CTR | 28 | 16 |
| **Tran** | 43811 | 3826 | CTR | 20 | 24 |
| **Franjic** | 187241 | 16342 | CTR | 37 | 37 |
| **Yang** | 52501 | 2003 | COVID-19 + CTR | 16 | 19 |
| **Fullard** | 22937 | 2142 | COVID-19 + CTR | 18 | 20 |

\*AD, Alzheimer's disease; CTR, healthy controls; LBD, Lewy Body Diseases; MS, Multiple sclerosis.

The clustering analysis of the 64,438 immune cell nuclei resulted in the identification of 15 clusters (Figure 1B), annotated 0 to 14 in descending order based on the total number of nuclei per cluster (Figure 1C). These clusters were distributed across all neurological and healthy settings (Supplementary Figure 4 and Supplementary Table 2). As expected, microglia were naturally the predominant cell type. In addition, we annotated clusters corresponding to other immune cell populations, including macrophages and T cells. Clusters 9 and 12 exhibited significant upregulation of markers associated with macrophages (*MRC1*, *CD163*) and T cells (*CD247*, *TRAC*, *CD3E* and *CD8A*), respectively, while showing minimal expression of microglial markers (*P2RY12*, *CX3CR1, AIF1, TMEM119*) (Figure 1D and 1E, and Supplementary Table 3). Two other small clusters, clusters 8 and 13, displayed high expression of markers associated with other CNS cell types. While cluster 8 appears to include doublets resultant from contaminant astrocytes (high expression of *SLC1A2* and *AQP4*), cluster 13 expressed high levels of markers of astrocytes, oligodendrocyte progenitor cells (OPCs) (*PDGFRA*) and neurons (*GRIK2*) (Figure 1D). Furthermore, these clusters presented a slightly increased number of mean genes per nuclei (Figure 1C). For this reason, clusters 8 and 13 were deemed doublet-containing clusters and were excluded from further analyses. The remaining 11 clusters (clusters 0-7, 10, 11 and 14) were identified as microglia subsets and had high expression of microglia markers (Figure 1D and 1E). Collectively, these microglia subpopulations encompassed 60,557 nuclei, composing the *HuMicA* integrated object.

Moreover, among the 11 microglia populations, clusters 3, 6 and 7 were analysed separately (Supplementary Figure 5A). These clusters had a high expression of markers associated with neurons (*SYT1*, *SNAP25*) and oligodendrocytes (*ST18, PLP1*), particularly in clusters 3 and 6, respectively (Supplementary Figure 5B and Supplementary Table 3). We considered cluster 7 to be closely related to clusters 3 and 6, as they shared similar expression patterns (Supplementary Figure 5B). Indeed, these subpopulations were not considered doublets since they successfully passed the strict pre-processing pipeline, which included doublet removal. Also, the number of mean genes per nuclei was proportional and the expression of microglia markers remained consistent. Regarding the distribution of these clusters in pathology, Cluster 7 showed an expansion in autism, while cluster 6 was expanded in LBD and MS, compared to controls (Supplementary Figure 5C). Notably, the behaviour observed in cluster 6 has been previously described and validated as a microglia subpopulation associated with excessive myelin phagocytosis, in a snRNA-seq study that is included in our integrated dataset^14^. Continuing with this line of reasoning, we can speculate that cluster 3 may indeed represent a subpopulation associated with excessive phagocytosis of synaptic terminals. Cluster 7, on the other hand, could possibly correspond to a preliminary state in relation to cluster 3. Interestingly, we observed an enrichment of cluster 6 in LBD and MS samples in comparison with the healthy population. Given that MS is a well-defined demyelinating syndrome, the observed results may shed light on the role of microglia in this neurological disorder, particularly with respect to myelin phagocytosis and its potential implications in disease progression.

### Homeostatic microglia clusters and their changes across neurodegenerative pathological conditions

The homeostatic microglia signature is primarily characterized by high expression of the so-called *homeostatic* genes, particularly *P2RY12* and *CX3CR1*. Several *HuMicA* clusters had the features of homeostatic microglia (Figure 2A). Clusters 0, 5 and 11 displayed increased expression of homeostatic genes (Figure 2B and 2C), further supported by the presence of *P2RY12* among their significantly upregulated differentially expressed genes (DEGs) (Supplementary Table 3). In addition, we also annotated cluster 10 as homeostatic, although the increased expression of homeostatic genes was not as pronounced as in the other clusters. Cluster 10 did not express high levels of any other specific marker (e.g., activation-related, inflammatory, etc) that could provide a more precise characterization. Moreover, the upregulated DEGs and related enriched gene ontology (GO) terms showed significant overlap with those observed in clusters 0, 5 and 11 (Supplementary Table 3 and Supplementary Figure 5D), further supporting its categorization as a homeostatic cluster. Cluster 4, on the other hand, expresses high levels of homeostatic genes. However, it also expresses high levels of markers associated with a pathological phenotype, such as *APOE* (Figure 2C), suggesting a pathology-associated nature to this cluster. This will be explained with greater detail below.

**Figure 2.**
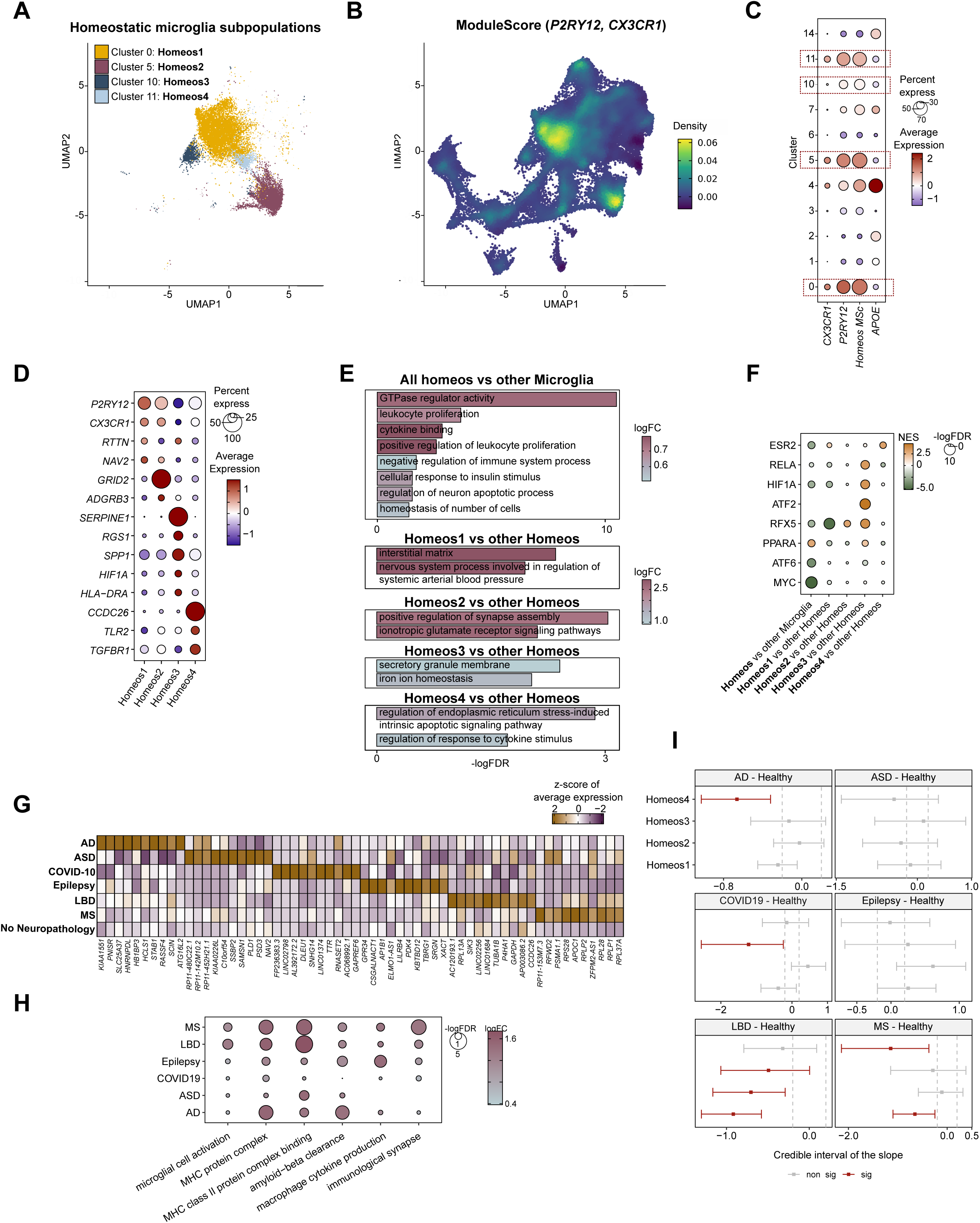
(A) UMAP representation of the four clusters annotated as homeostatic microglia. Cluster 0, *Homeos1*; Cluster 5, *Homeos2*; Cluster 10, *Homeos3;* Cluster 11, *Homeos4*. **(B)** UMAPs showing the density of the expression of the module score (MSc) of homeostatic markers (*P2R12* and *CX3CR1*) in the *HuMicA*. **(C)** Dot plot representation of the expression of *P2RY12, CX3CR1, APOE,* and Module Score (MSc) expression of homeostatic markers. **(D)** Dot plot representation of the expression of genes upregulated in all homeostatic clusters vs other microglia and specifically in each of the homeostatic subpopulations. **(E)** Representation of selected GO terms enriched within upregulated DEGs in homeostatic clusters vs other microglia and specifically in each of the homeostatic subpopulations. Enrichment is represented by the negative of the log of the adjusted p value (FDR) and the log of the fold change. **(F)** Representation of the enrichment of a selected group of TFs relative to the comparison of differential expression of all homeostatic clusters vs all other microglia, and specially in each of the homeostatic subpopulations. Significance is represented by the Normalized Enrichment Score (NES) and the negative of log of the adjusted p value (FDR). **(G)** Heatmap representation of the z-score of the average expression (by pathological condition) of the top ten most upregulated genes in homeostatic microglia that are exclusive to each pathology when compared with the healthy group. **(H)** GO enrichment of selected categories obtained from the lists of upregulated DEGs in homeostatic nuclei in each pathology vs controls. Enrichment is represented by the negative of the log of the adjusted p value (FDR) and the log of the fold change. **(I)** Differential distribution of each homeostatic clusters in each pathology in relation to the healthy group. Population expansion or depletion is represented by the credible interval of the slop. Statistical significance is considered for an adjusted p value (FDR) < 0.05 and is highlighted in red.

To describe both general homeostatic and subpopulation-specific traits, we obtained DEGs by comparing all homeostatic nuclei vs all other microglia clusters and comparing each homeostatic subpopulation with the remaining three (Supplementary Table 3). The GO enrichment analysis of the upregulated DEGs in all homeostatic clusters vs all other microglia cells showed the highest significance for terms involved in GTPase activity. Additionally, it yielded categories such as *leukocyte proliferation*, *negative regulation of immune system process* and *homeostasis of number of cells* (Figure 2E and Supplementary Figure 6B). Among the homeostatic clusters, *Homeos1* (cluster 0) appeared to represent the “classic” homeostatic microglia phenotype, displaying the highest levels of *P2RY12* and *CX3CR1*, and it also shows specific upregulation of *RTTN* and *NAV2* (Figure 2D and Supplementary Figure 6A). *Homeos2* (cluster 5) showed high expression levels of genes such as *GRID2* and *ADGRB3* (Figure 2D and Supplementary Figure 6A), which are associated with GO terms related to *synapse assembly* (Figure 2E and Supplementary Figure 6B). The subpopulations *Homeos3* (cluster 10) and *Homeos4* (cluster 11) exhibited lower expression of homeostatic markers and upregulated activation-related genes compared to the other homeostatic clusters. *Homeos3* showed upregulation of *SERPINE1*, *SPP1* (classic DAM gene), *HIF1A* and *HLA-DR*; and *Homeos4* displayed increased expression of more typical inflammatory signalling genes like *TLR2* and *TGFBR1* (Figure 2A and Supplementary Figure 6A). These data suggest that *Homeos3* and *Homeos4* subpopulations might correspond to preliminary stages of the homeostatic-to-activated shift in microglia.

To gain insights into the potential transcription factor (TF) involvement, we utilized gene expression data of their targets for all the comparisons mentioned above. By employing the DoRothEA tool, we predicted the activity of regulatory TFs, based on the differential expression common to all homeostatic clusters compared to other microglia, as well as ones particular to each homeostatic subpopulation. MYC and PPARA present decreased and increased association, respectively, in homeostatic microglia compared to all other microglia. RFX TFs, known activators of MHC-II and linked to autism and cognitive disabilities^25^, showed a lower involvement in *Homeos1* and, in opposition, have a higher association with *Homeos2* and *Homes3*. The latter showed specific involvement of TFs associated with inflammatory activation such as HIF1A and RELA (Figure 2F and Supplementary Figure 6C).

To explore the potential differences in homeostatic microglia across different pathologies, we calculated DEGs on the bulk of homeostatic nuclei in each pathology compared to healthy controls (Supplementary Table 4). While some genes showed similar behaviour across multiple pathologies, many genes showed either upregulation or downregulation exclusively in a specific pathology in comparison to the healthy population (Figure 2G and Supplementary Figure 6D). We observed that homeostatic microglia from neurodegenerative patients displayed upregulation of genes associated with microglia activation and pro-inflammation. This was also evidenced by the GO terms enriched within each list of upregulated DEGs in each pathology, which included categories associated with antigen presentation, amyloid-beta clearance, and cytokine production. Of note, the only pathology that did not show this primed state of homeostatic microglia was COVID-19, the only non-standard neurologic disease (Figure 2H).

To gain further insights into potential associations with neurodegenerative pathologies, we examined the distribution of the bulk of the homeostatic microglia cells and each homeostatic cluster individually across the pathologies included in the *HuMicA*. Our analysis revealed varying patterns of behaviour across different conditions. However, a notable trend emerged, indicating a depletion of the homeostatic subpopulations in four out of six pathologies (AD, COVID-19, LBD and MS) for at least one homeostatic cluster compared to the healthy population (Figure 2I and Supplementary Figure 6E and 6F).

### The spectrum within the disease-associated microglia (DAM) phenotype in neurodegenerative conditions

The occurrence of a DAM gene signature has been the gold standard for characterizing microglia activation^5^. We measured the module score (MSc) expression of the full list of human genes in the DAM signature (n=210), as previously reported by Thrupp *et al.* (2020)^26^. Our observations revealed a notable concentration of this signature within three microglia subpopulations, clusters 1, 2 and 4 (Figure 3B and 3D). In a recent study, Silvin and colleagues, described in mice two DAM populations with distinct ontogeny^27^. One population corresponds to legitimate DAM, while the other consists of disease-inflammatory macrophages (DIMs), which represent a microglia-like population derived from infiltrating monocytes. Using the summarized human gene signatures provided in the aforementioned study, we found that cluster 2 specifically overexpresses DIM genes. Notably six genes from the original DIM signature were present among the upregulated DEGs of the DIM in cluster 2 when compared to DAMs in clusters 1 and 4 (*CD83, FOS, BTG2, SAT1, CSF2RA, ZFP36*) (Figure 3C and 3D, Supplementary Figure 7A and Supplementary Table 3). The *DIM* phenotype observed in cluster 2 is supported by the enrichment of multiple pro-inflammatory pathways associated with interleukin signalling, which distinguishes from the two DAM clusters (Figure 3E and Supplementary Figure 7B and 7C). To further investigate the characteristics of DAM activation, we performed a comparison between cluster 1 and cluster 4. Interestingly, these two clusters showed clear segregation based on the expression of *SPP1* and *APOE*, which were specifically increased in cluster 1 and cluster 4, respectively (Figure 3C). These findings align with the two stages of DAM activation previously described by Keren-Shaul *et al.* (2017) and referred to as stage 1 DAM (intermediate) and stage 2 DAM (final activation)^5^. Employing the gene sets associated with these DAM stages, we annotated cluster 4 as the *Intermediate.DAM* and cluster 1 as the *Final.DAM*. The enrichment of the stage 1 phenotype is particularly evident in cluster 4 (Supplementary Figure 8A), where four out of seven genes from the original gene set are upregulated in the *Intermediate.DAM* when compared to the *Final.DAM* comparison (*APOE, FTH1, B2M, TYROBP*) (Figure 3F). The demonstration of the stage 2 phenotype in the *Final.DAM* (cluster 1) was not as straightforward when considering the original signature (Supplementary Figure 8A). Only *ITGAX,* among the genes from original signature, was found to be upregulated in the *Final.DAM* compared to *Intermediate.DAM* (Figure 1F). Nevertheless, GO of the upregulated DEGs when comparing *Final.DAM* and *Intermediate.DAM* revealed a strong enrichment of terms associated with phagocytosis (Figure 3G and Supplementary Figure 8B), which is consistent with the characteristics originally described^5^.

**Figure 3.**
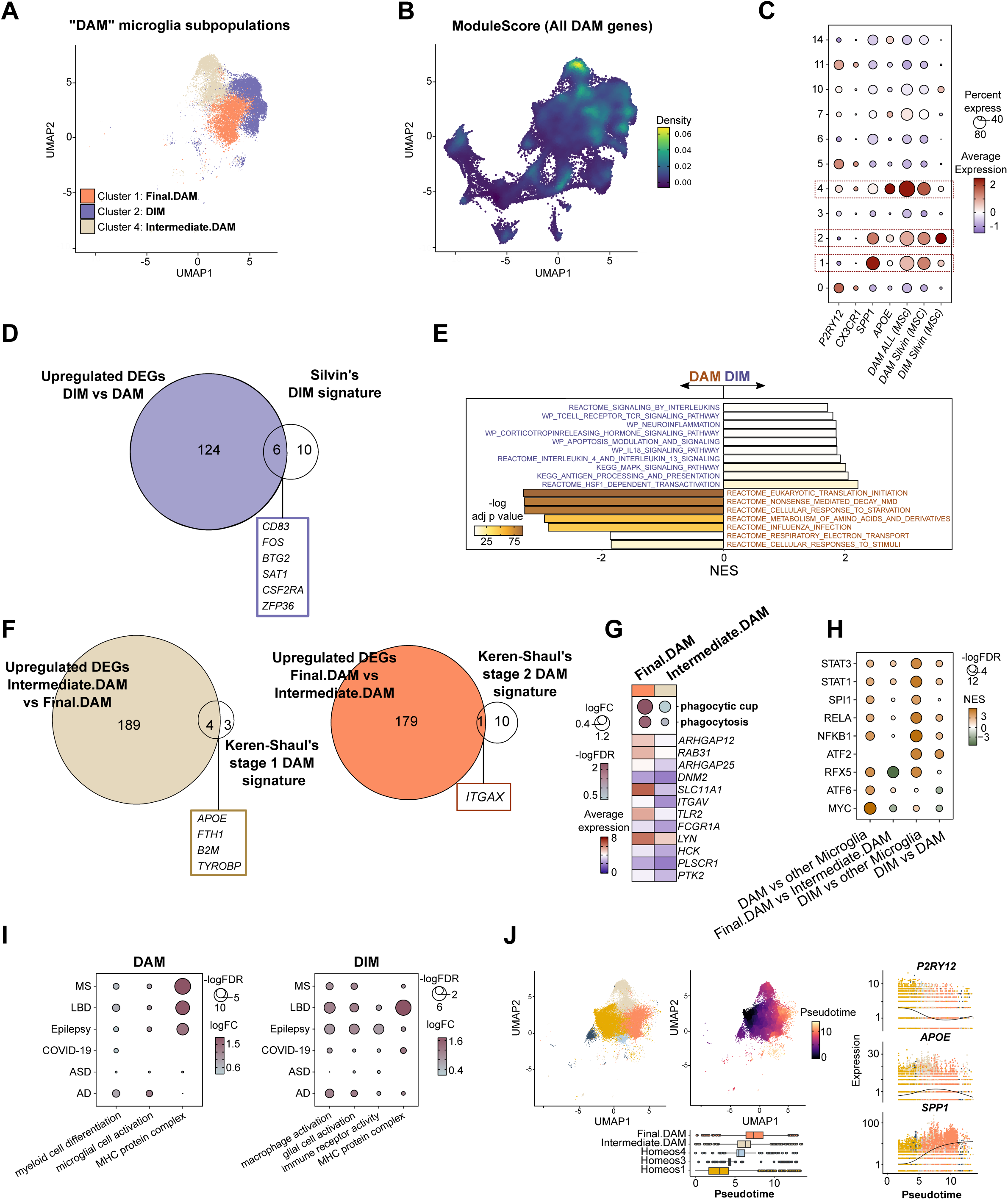
(A) UMAP representation of the three clusters considered within the “DAM universe”. Cluster 1, *Final.DAM*; Cluster 2, *DIM*; Cluster 4, *Intermediate.DAM*. **(B)** UMAPs showing the density of expression of the module score (MSc) of all 210 genes from the human DAM phenotype in the *HuMicA*. **(C)** Dot plot representation of the expression of *P2RY12, CX3CR1, SPP1, APOE,* and MSc expression of all DAM genes, the conserved DAM human signature (*SPP1, B2M, TYROBP, TREM2, MAMDC2, CD63, GPNMB, CCL3*) and the conserved DIM human signature (*CCL4, CD14, CD83, CSF2RA, EIF1, FOS, IER2, JUN, JUNB, EGR1, IL1B, TNF, PLAUR, SAT1, ZFP36, BTG2*) from Silvin *et al.* (2022)^27^ **(D)** Venn diagram showing the overlap between the genes significantly upregulated in our *DIM* vs *DAM* comparison and the conserved DIM human signature. **(E)** Barplot representing selected GSEA enriched terms from the *REACTOME, WILKIPATHWAYS* and *KEGG* repositories for the differential expression of *DIM* vs *DAM* subpopulations. Enrichment is represented as a function of the negative of the log of the adjusted p value and the Normalized Enrichment Score (NES). **(F)** Venn diagram showing the overlap between the genes significantly upregulated in our *Intermediate.DAM* vs *Final.DAM* comparison and a set of genes characteristic of stage 1 DAM phenotype (*APOE, B2M, TYROBP, FTH1, LYZ, CTSB, CTSD*); and the genes significantly upregulated in our *Final.DAM* vs *Intermediate.DAM* comparison and a set of genes characteristic of stage 2 DAM phenotype (*LPL, CTSL, TREM2, AXL, CD9, CSF1, CCL23, ITGAX, CLEC7A, LILRB4, TIMP2*). Both stage 1 and stage 2 gene signatures were developed based on Keren-Shaul *et al* (2017)^5^. **(G)** GO enrichment of selected categories associated with phagocytosis obtained from the lists of upregulated DEGs in *Final.DAM* vs *Intermediate.DAM*, coupled with average expression (by cluster) of the genes contributing for the enrichment. Enrichment is represented by the negative of the log of the adjusted p value (FDR) and the log of the fold change. **(H)** Representation of the enrichment of a selected group of TFs relative to the comparison of differential expression of *DAM* vs other microglia, *Final.DAM* vs *Intermediate.DAM*, *DIM* vs other microglia and *DIM* vs *DAM*. Significance is represented by the NES and the negative of log of the adjusted p value (FDR). **(I)** GO enrichment of selected categories obtained from the lists of upregulated DEGs in DAM and DIM nuclei in each pathology vs controls. Enrichment is represented by the negative of the log of the adjusted p value (FDR) and the log of the fold change. **(J)** Trajectory analysis in a subset Seurat object composed by the standard homeostatic microglia (*Homeos1*, cluster 0), the pre-activated microglia (*Homeos3*, cluster 10; *Homeos4*, cluster 11), and the DAM populations (*Intermediate.DAM,* cluster 4; *Final.DAM*, cluster 1). *Homeos1* was used as starting point. The nuclei represented in the UMAPs are annotated by each of the corresponding cluster and by the pseudotime. The boxplot represents the distribution of all nuclei grouped by cluster across the pseudotime. The expression values of *P2RY12, APOE* and *SPP1* are represented in relation to the pseudotime and grouped by cluster.

We also evaluated the specific TF enrichment for both DAM and DIM subpopulations (Figure 3H and Supplementary Figure 9A). In contrast to what we described for homeostatic microglia, the DAM nuclei showed high association with MYC and ATF6, both of which have been previously described as important factors in microglia activation^28,29^. Additionally, increased involvement of RFX TFs appeared to be characteristic of the *Intermediate.DAM* and DIMs. Moreover, DIMs exhibited enhanced involvement of ATF2, PU.1, NF-kB subunits, STAT1 and STAT3, supporting their overall pro-inflammatory profile.

We also examined the differential expression between each pathology and the control group for the bulk of DAM populations and for the DIMs (Supplementary Figure 9B and Supplementary Table 4). Similar to what was observed for homeostatic microglia, we identified lists of DEGs exclusive for each of the diseases. In both DAMs and DIMs from patients, there was an upregulation of genes associated with a more primed state characterized by a high enrichment of GO terms linked to microglia activation and antigen presentation. However, the extent of enrichment varied across different pathologies, and once again, this trend was not observed for COVID-19 (Figure 3I).

Regarding the distribution of each of these three clusters across pathological conditions (Supplementary Figure 9C and 9D), there was no consistent and statistically significant differential distribution in relation to the healthy population. The only exceptions were observed as an expansion of *Intermediate.DAM* in autism and a depletion of *DIM* in LBD. However, it is essential to consider the heterogeneity of the distribution of subjects in each group, especially since samples from ASD and LBD come from only one dataset each.

To gain further insights into the landscape of the DAM activation process, we conducted a trajectory analysis on a subset Seurat object comprising nuclei from standard homeostatic microglia (*Homeos1*, cluster 0), pre-activated microglia (*Homeos3*, cluster 10; *Homeos4*, cluster 11), and the DAM populations *(Intermediate.DAM*, cluster 4; *Final.DAM*, cluster 1). The analysis started at *Homeos1* as the initial point. The pre-activation homeostatic clusters (*Homeos3* and *Homeos4*) were positioned at the mid-point of the trajectory, while both DAM clusters were located at the end of the trajectory. Interestingly, the *Intermediate.DAM* cluster appeared slightly left-shifted in the pseudotime relative to the *Final.DAM*, which further validates the annotation of these clusters. Critical genes involved in the stage 1 and stage 2 DAM activation displayed significant differential expression across the trajectory. Notably, there was a decrease in *P2RY12* expression from homeostatic-to-DAM transition, while *APOE* and *SPP1* exhibited opposite patterns of behavior. *APOE* showed early upregulation, whereas *SPP1* showed upregulation more prominently in later stages of activation (Figure 3J and Supplementary Table 5).

### Changes in DAM subpopulations across neurodegenerative conditions

We aimed to characterize more precise subpopulations or substates within the *Final.DAM, Intermediate.DAM* and *DIM* populations. To achieve this, each subpopulation as further split into four smaller substates (Figure 5A). We found that each subpopulation showed distinct differential expression patterns relative to each other (Supplementary Figure 10A and Supplementary Table 3). Interestingly, these niche subpopulations demonstrated statistically significant differences in distribution between patients and healthy controls. Most markedly, we observed an expansion of *Final.DAM.3* in AD and MS, *DIM.1* and *DIM.2* in epilepsy, and *Intermediate.DAM.3* in ASD and MS (Figure 4B and Supplementary Figure 10B). Further exploration of the upregulated DEGs and the corresponding GO terms for each of these subpopulations revealed distinct functional characteristics. *Final.DAM.3* have upregulated expression of genes associated with ameoboidal cell migration; *Intermediate.DAM3* exhibited a more pro-inflammatory pattern associate to higher cytokine production; *DIM.3* was linked to myeloid differentiation; and *DIM.1* showed an increased expression of genes involved in the response to TGF-β (transforming growth factor beta) (Figure 4C and Supplementary Figure 10C).

**Figure 4.**
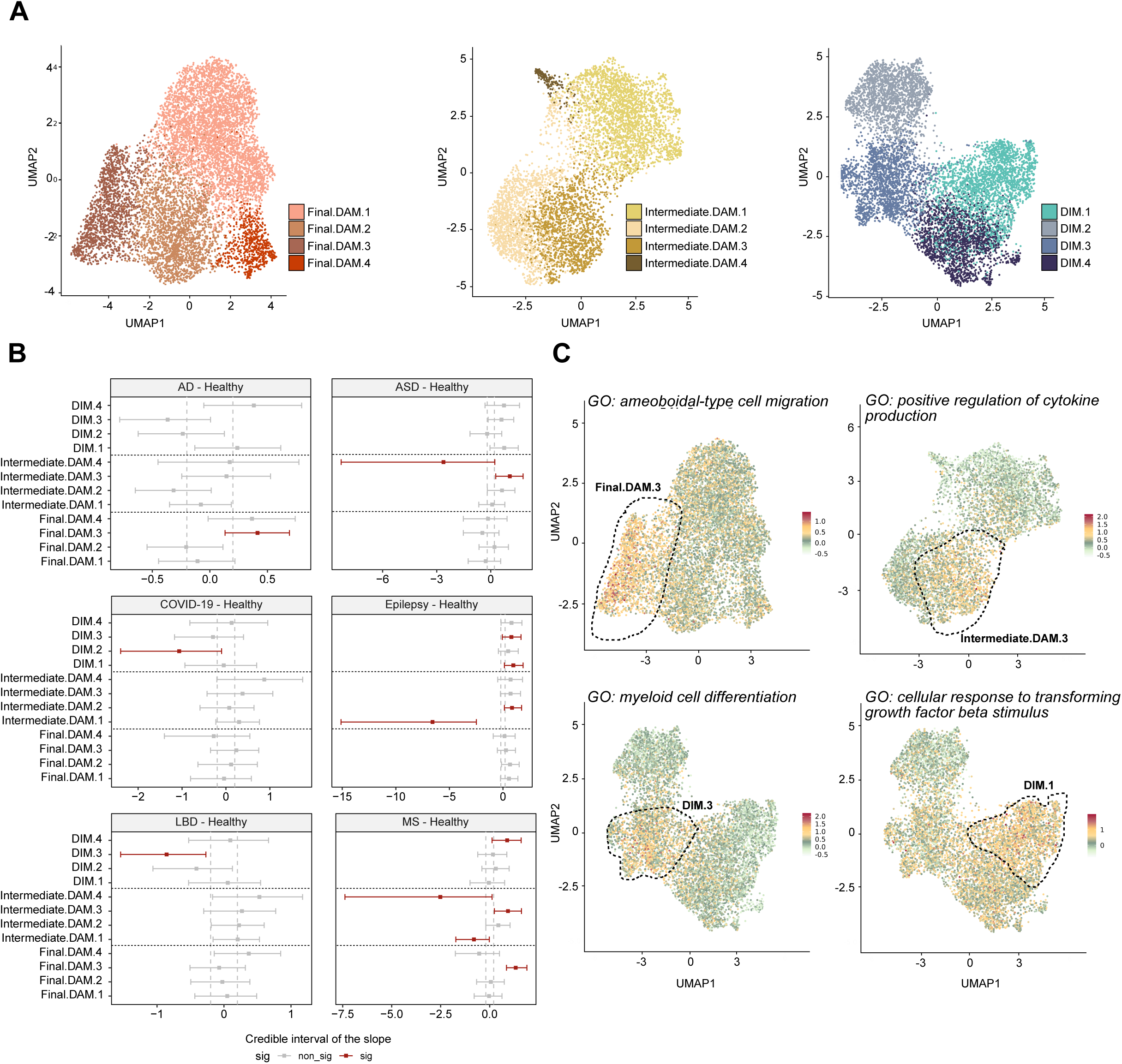
(A) UMAP representation of the four subpopulations annotated within the *Final.DAM, Intermediate.DAM* and *DIM* clusters. **(B)** Differential distribution of each subpopulation in each pathology in relation to the healthy group. Population expansion or depletion is represented by the credible interval of the slop. Statistical significance is considered for an adjusted p value (FDR) < 0.05 and is highlighted in red. **(C)** UMAPs showing the module score (MSc) expression the sets of upregulated genes in each subpopulation associated with the corresponding enriched gene ontology (GO) terms (e.g., the *ameoboidal-type cell migration* GO term is enriched for the upregulated DEGs in *Final.DAM.3* vs other *Final.DAM*).

**Figure 5.**
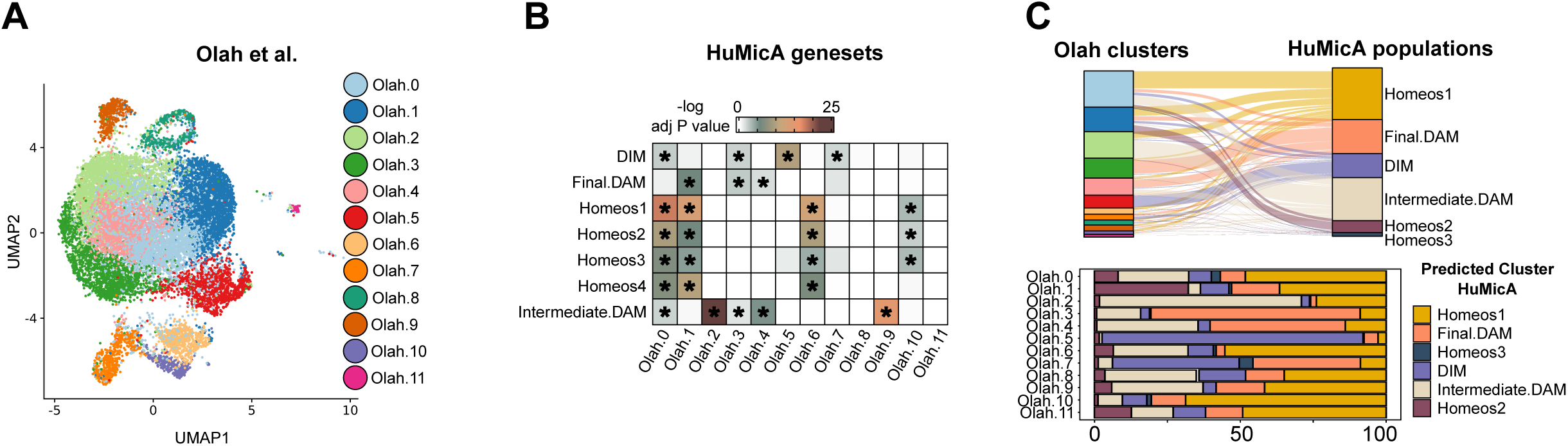
(A) UMAP visualization of the scRNA-seq dataset from Olah et al. (2020) consisting of 16,130 sorted microglia cells, clustered in twelve subpopulations. **(B)** Heatmap representation of the GSEA enrichment of the lists of genes upregulated in the *HuMicA* subpopulations for the differential expression of each *Olah* cluster. Enrichment is represented as a function of the negative of the log of the adjusted p. The asterisk represents the enrichments with adj p value < 0.05. **(C)** Sankey plot depicting the estimated prediction of association of the *Olah* cells in relation to the microglia populations in the *HuMicA*. Each line connects each cell of the *Olah* clusters with the *HuMicA* subpopulation showing the maximal prediction score. Complementary stacked bar plot representation of the percentage of cells from each *Olah* cluster that show the highest prediction to the respective *HuMicA* clusters.

### External validation in an independent dataset

To validate the signatures described in the *HuMicA,* we leveraged a publicly available scRNA-seq dataset published by Olah *et al.* (2020)^30^. This dataset comprises immune cells isolated from a mix of autopsied dorsolateral prefrontal cortex obtained from both AD patients and controls with mild cognitive impairment, as well as surgically resected cortex samples near the epilepsy focus of temporal lobe epilepsy patients. After filtering out non-microglial subpopulations, we obtained a total of 16,130 cells, which were clustered in 12 subpopulations (Figure 5A). To assess the enrichment of the signature genesets from the *HuMicA*, we conducted Gene Set Enrichment Analysis (GSEA) on the lists of upregulated genes in the subpopulations of our study, using the differential expression calculated for each of the *Olah* clusters (Figure 5B). We found that the genesets from the four homeostatic clusters are enriched in a relatively indiscriminate manner in Olah.0, Olah.1, Olah.6 and Olah.10. While the bulk homeostatic profile appears to be replicable, the subpopulation-specific traits appear not to be as precise. On the other hand, the microglia populations from the “DAM universe” displayed enrichment in specific and distinguished *Olah* clusters. The genes upregulated in *DIM* were highly enriched exclusively in *Olah.5* and *Olah.7*, while the *Intermediate.DAM* signature was overrepresented in *Olah.2* and *Olah.9*. Regarding the *Final.DAM* geneset, no clear enrichment was observed. Although some Olah clusters are significantly enriched for it, it is not exclusive. To further investigate the prevalence of the *HuMiCa* signatures and to clarify the mixed enrichment observed inthe previous analysis, we performed a prediction of the probability of each cell from *Olah* annotating to the clusters of the *HuMicA* (Figure 5C and 5D). The patterns observed in this analysis were consistent with the those identified in the GSEA analysis and complemented it. The *Homeos1* cluster showed the highest prediction score for the majority of cells of *Olah.0*, *Olah.6, Olah.10* and also *Olah.11*. Notably, Olah.1 exhibited mixed prevalence of *Homeos1* and *Homeos2*, highlighting the challenge in discriminating between the homeostatic states. The *Intermediate.DAM* phenotype was more prevalent in *Olah.2*. Cells from *Olah.3* and *Olah.4* showed strong relationship to the *Final.DAM,* and *Olah.5* associated with *DIM*. No high prediction was observed for the smaller pre-activation homeostatic populations from the HuMicA, *Homeos3* and *Homeos4*. The distinctive enrichment of each *HuMicA* geneset within specific Olah clusters demonstrates that the populations observed in our integrated analysis are prevalent in external and independent data, reinforcing the biological replicability of the findings.

## DISCUSSION

Our integrated analysis of snRNA-seq datasets comprising brain biopsies from various neurological pathologies and healthy subjects, has resulted in the creation of the first comprehensive human atlas of microglia called *HuMicA*, which includes 60,557 microglial nuclei. The clustering analysis of the integrated object confirmed that microglia encompass a range of distinct cell subpopulations rather than a one-directional transition from homeostasis to activation. In mice, several studies have already developed atlases covering the spectrum of microglia throughout embryonic development, aging, and in pathological conditions^27,31,32^. The translation of these findings into human biology has become feasible due to the availability of snRNA-seq studies, which have significantly contributed to the more precise characterization of the human cerebral environment, by identifying specific cellular subpopulations. However, microglia often take on secondary attention in human studies primarily due to their low abundance within bulk brain samples. In our analysis, the average proportion of the immune cell population relative to the entire dataset was approximately 5%, which aligns with the reported amounts in the original studies. This limitation has previously impacted the scalability of the studies, requiring larger number of samples, and hinders the identification of smaller microglia subpopulations, which remain underrepresented in smaller datasets. In this study, we have successfully overcome this significant limitation by employing a bioinformatic strategy that maximizes the utilization of already published data.

The *HuMicA* comprises eleven microglia subpopulations, with ten of them containing a substantial number of cells, allowing for detailed characterization. For instance, we successfully described the classical homeostatic microglia phenotype, which accounted for the most abundant population, *Homeos1*. In addition, we identified three additional populations with a homeostatic gene profile. Two of these subpopulations were smaller in size and likely represent two distinct stages of pre-activation, one leaning towards DAM activation (*Homeos3*) and the other tending towards general pro-inflammation (*Homeos4*). Special attention must also be given to *Homeos2*, which expresses high levels of *P2RY12* and *CX3CR1*, and we annotated as “homeostatic”. Indeed, *Homeos2* presents a distinct profile specifically driven by high expression of *GRID2* and *ADGRB3*, which segregates it clearly from the other subpopulations Interestingly, a similar microglia population, denominated AD2 was identified in an external snRNA-seq study. This AD2 subpopulation was found to be expanded in AD compared to controls and was characterized by upregulation of *P2RY12, CX3CR1, GRID2* and *ADGRB3.* Additionally, this gene expression pattern was also associated with synapse assembly and specifically related to a response to tau-bearing dying neurons^33^. The true nature of the *Homeos2* is subpopulation still requires further studies to elucidate whether it behaves as a homeostatic microglia subpopulation or represents a distinct pathology-related state.

Previous studies had already demonstrated an association between microglia activation and neuropathology. In these, the typical strategy often involves assessing differential gene expression in microglia in from patients compared to healthy controls^7–12,14,16,19,20^, with a general description of a pathology-related pro-inflammatory and activated state. In addition, some studies have shown a differential distribution of microglia subpopulations between patients and controls, with the transcriptomic profile of these subpopulations enriched in genes associated with activation. This includes microglia subpopulation resembling the DAM phenotype observed in mice, as well as those expressing genes and in genes associated with genetic risk factors for neurological diseases^7,8,11,15^. The DAM phenotype has indeed been a valuable tool in evaluating the role of microglia in neurodegeneration. It has been shown to be prevalent not only in AD mice, but also in a model of amyotrophic lateral sclerosis^5^. Furthermore, the aforementioned DAM enrichment in pathology-related microglia extend beyond AD^7,8,11^, encompassing other conditions such as autism^15^ and severe COVID-19^19^, emphasizing the inter-disease and unspecific nature of DAM. Indeed, the role of the DNA phenotype in neuropathology, particularly in human AD, has been a topic of debate, and its replicability has been questioned^12^. Moreover, from the methodological standpoint, it has been shown that snRNA-seq microglia express lower levels of main DAM genes in comparison to scRNA-seq microglia^26^. However, in the case of *HuMicA*, we exclusively used snRNA-seq data and were still able to widely demonstrate detectable and varying levels of DAM genes across the microglia populations of our datasets.

It is reasonable to conclude that the DAM phenotype alone may not be sufficient to fully understand microglia’s behaviour in pathology. The original study proposed a sequential two-stage process for DAM activation, with the first stage being *Trem2*-dependent stage 1 and the second stage being *Trem2-*independent^5^. However, a recent groundbreaking study by Silvin and colleagues (2022) provided further insights “deconstructing” the DAM signature. These authors developed an encompassing myeloid cell map of mice brain samples by integrating scRNA-seq data from different neuropathological settings. Within this “M-verse”, they observed two DAM populations with different ontogenies and concluded that one of these populations is composed by infiltrating monocyte-derive “disease-inflammatory macrophages” (DIMs). The authors demonstrated that *bona fide* DAMs were *Trem2*-dependent, while DIMs were *Trem2*-independent^27^. However, the authors did not elaborate on whether the DIMs could represent what was initially described as the stage 1 DAM based on the dual profile of *Trem2*-dependency. Here, we have shed light on that matter by describing the simultaneous prevalence of all these phenotypes, including the *bona fide* DAM, with the stage 1 DAM (*Intermediate.DAM*) and stage 2 phenotype (*Final.DAM*), and the *DIM* population. It is worth noting that the DIM population clustered closely with the macrophage population in the *HuMicA*. Regarding the *Trem2-*dependency, our experimental design did not allow for its direct evaluation, and we observed the expression of across all *TREM2* populations.

An important aspect used to interpret distinct microglia subpopulations is the differential expansion or depletion in different pathological conditions. We observed a decrease in homeostatic clusters in a generalized way across the studied diseases. As for the DAM and DIM subpopulations, we noticed altered distribution of smaller and more precise subpopulations within the *Final.DAM, Intermediate.DAM* and *DIM* clusters, with differential patterns across pathologies. For example, we described two DIM subpopulations, associated with differentiation, cytokine binding and response to TGF-B, which were expanded exclusively in epilepsy. In addition, a *Final.DAM* subpopulation with migration-related phenotype is increased in AD and MS, while an *Intermediate.DAM* subpopulation was more prevalent in patients with ASD and MS. DAM populations have consistently been shown to be expanded in pathology, particularly in AD. In our study, we were able to identify niche substates within this spectrum with distinct profiles differentially distributed in pathology, rather than observing a general expansion of the DAM phenotype. We must consider the potential confounding effect caused by over-normalization of the data, which is inherent to the integration of multiple datasets. Therefore, subsequent validations are required to evaluate whether these associations between pathologies and populations are genuine and not random outputs of statistical bias.

The *HuMicA* cluster 6 showed a clear and challenging-to-refute expansion in MS and LBD. This subpopulation exhibited detectable levels of oligodendrocyte markers, such as *ST18, PLP1,* and *MBP*. Interestingly, this pattern has been previously observed and validated *in vitro* in a study whose data we included in the *HuMicA.* In that study, the authors incubated human primary microglia with myelin particles and observed the localization of oligodendrocyte-derived mRNA within the phagocyting microglia and close to the nuclei^14^. Similar patterns of phagocytic microglia engulfing oligodendrocyte-derived mRNA were described in mice scRNA-seq data^32^. This suggests that cluster 6 in the *HuMicA* could potentially be related to microglia ingulfing oligodendrocyte-derived material. By the same rationale, one could speculate that clusters 3 and 7 of the *HuMicA* may be associated with excessive pruning of synaptic terminals. However, we decided not to highlight these subpopulations in our subsequent analyses due to the potential implications of non-microglial RNA presence within the microglia transcriptomes impact the interpretation of the data. Nevertheless, the demyelinating nature of MS raises questions on the biological significance of these results.

The findings described in any integrated object must be considered within the limitations intrinsically related to *in silico* integration of multi-source data, which may potentially mask or exaggerate biologically relevant traits. We acknowledge the necessity for validation of our findings through additional approaches. *In situ* hybridization and immunohistochemistry of brain tissue slides would be valuable in corroborating the prevalence and coexistence of these microglia subpopulations, and further elucidating their potential pathology-related functions. Nonetheless, we successfully validated *HuMicA* in an independent scRNA-seq dataset, which was published as an atlas of human microglia subpopulations on its own^30^. We observed that the transcriptomic profiles characteristic of the main subpopulations in the *HuMicA* were indeed associated with specific microglia clusters in the external dataset. This approach not only validated the presence of the *HuMicA* patterns but also revealed a “clean” translation without major overlapping of its signatures on the clusters of the external data.

Using the *in-silico* approach in our study, we successfully compensated for the common limitation of low sample size in individual human microglia snRNA-seq datasets, which have significantly prevented the ability to fully characterize microglial subpopulations. We have described and validated the *HuMicA,* in which we describe multiple homeostatic microglia states as well as pathology-related phenotypes, only described in animal models up to now. We intend that the *HuMicA* may serve as a public resource to study microglia under multiple experimental settings thus contributing for the evolution of the overall knowledge on microglial biology.

## METHODS

### Single nucleus (sn)RNA-seq datasets

The human brain tissue snRNA-seq data used in this study was obtained from 16 public datasets (Table 1). Detailed technical and clinicodemographic characterization of the used samples within each dataset is presented in Supplementary Table 1. We included six datasets for AD^7–12^, two for MS^13,14^, two for epilepsy^26,34^, two for COVID-19^19,20^, one each for LBD (including Parkinson’s, Parkinson’s with dementia and dementia with Lewy bodies)^16^ and autism spectrum disorder^15^. The control groups within each study were included in the “No Neuropathology” group, plus data extracted from two additional datasets exclusively with individuals with no neuropathological alterations^24,35^. Only unsorted snRNA-seq data was included in the study. After pre-processing, 295 individual samples contributed to the atlas.

### Dataset pre-processing

A dataset downloaded as raw fastq files (Jakel) was processed using the CellRanger v.6.1.2 software^36^. Individual count data was obtained with the *count* function, aligning to the reference genome GRCh38-3.0.0. The *--include-introns* option was used to map both unspliced pre-mRNA and mature RNA. Individual count matrices were generated for each library and concatenated into a single count matrix.

Most datasets were obtained already in the count matrix format, either at the raw or pre-processed stages. Data had been already demultiplexed and sequences aligned to the human reference genome (GRCh38), accounting for both intronic and exonic regions. For each dataset, count data from each subject was concatenated in one single count matrix. Data processing and subsequent analysis was performed using the *Seurat* (v4.0.2) R package^37^. Quality control filtering was uniformized to all datasets. Genes were only included if detected in at least 3 nuclei. Nuclei with unique genes inferior to 200 or superior to 5,000, total UMI counts less than 500 or over 20,000, mitochondrial RNA content superior to 20% or ribosomal RNA content superior to 5% were filtered out. Furthermore, a list of 105 genes described as related to postmortem interval in cerebral cortex was excluded^38^. The *doubletFinder_v3* function^39^ was used to estimate doublets from each individual subject, which were removed from each dataset’s Seurat object, followed by *SCTransform* normalization^40^ with default 3000 variable genes. Dimensionality reduction was performed with Principal Component Analysis (PCA) followed by Uniform Manifold Approximation and Projection (UMAP) accounting for the 30 main principal components (PCs).

### Cell-type annotation and isolation of immune cells

Cell clustering was performed with *FindNeighbors* (30 PCs) and *FindClusters* (res = 0.05-0.08) functions. The low resolution used allowed for a broad characterization of the 6 main cell types in brain tissue: neurons, oligodendrocytes, OPCs, astrocytes, endothelial cells, and immune cells (Supplementary Figure 1). The immune cell clusters encompassed microglia and other infiltrating or resident immune cells, such as T cells and macrophages. Cell type annotation was based on the identification of known canonical gene markers within the lists of clusters differentially expressed genes (DEGs) obtained using *FindAllMarkers* (*min.pct = 0.1 and logfc.threshold = 0.25, test.use=Wilcox*). *CD74, DOCK8, APBB1IP, HLA-DRA, PTPRC, P2RY12, C1QB, CX3CR1, C3, CSF1R* and *AIF1* were considered for identification of the immune cell clusters. The annotated clusters of immune cells did not present any of the known canonical genes for other brain cell types among its markers. Individual subjects with markedly low nuclei number in relation to the remaining dataset were removed before integration.

### Integration and clustering of the Human Microglia Atlas (HuMicA)

The immune cells from all datasets were integrated using the Seurat integration pipeline^41^ adapted for datasets normalized with *SCTransform*. The *FindIntegrationAnchors* and *IntegrateData* functions considered the 20 main PCs and the 3000 most variable features across all datasets. Dimensionality reduction was performed using PCA and UMAP (50 PCs). Clustering was performed using *FindNeighbors* (50 PCs) and *FindClusters* (res = 0.25), which outputted fifteen subclusters from a total of 64,438 nuclei (Figure 1).

### Inference of covariate influence

The integrated Seurat object was converted using the as.*SingleCellExperiment*. The percentage of variance explained, considering the genes in the “integrated” assay, was calculated using the *plotExplanatoryVariables* function from the *scater* package^42^ for the “Subject”, “Study”, “Group”, “Tissue region”, “Tissue state”, “Gender” and “Age” variables. Of note, the Fullard dataset does not supply gender nor age information, thus the corresponding samples were annotated as “NA”.

### Cluster characterization, calculation of DEGs and enrichment analysis

For the calculation of the DEGs, the “RNA” assay from the integrated object was normalized and scaled using the *NormalizeData* and *ScaleData* functions with standard settings. DEGs were calculate with the functions *FindAllMarkers* or *FindMarkers* (*min.pct = 0.25 and logfc.threshold = 0.25, test.use=MAST*). DEGs were considered significant if FDR < 0.05. For the visualization of the expression of specific genes or sets of genes, we used the normalized counts of the “RNA” assay, as abovementioned. The average expression of sets of multiple genes was obtained with *AddModuleScore*. Representation of gene expression in the UMAPs was accomplished with the *FeaturePlot* (Seurat) and *plot_density* (*Nebulosa*^43^) functions. Gene ontology (GO) enrichment was obtained using the *enrichGO* function from the *clusterProfiler* package^44^. For the netplots, the *simplify* function from *enrichGO* was used to remove redundancy of the most significantly enriched terms. The *dorothea*^45^ and *viper*^46^ packages were used to assess TF activity, accounting for the A regulon. The *fgsea* package^47^ was used to perform GSEA from differential expression comparisons. For both TF enrichment and GSEA, the results from *FindMarkers* (*min.pct = 0.1 and logfc.threshold = 0, test.use=MAST*) were used as input, and the avg_log2FC and the p_val_adj settings were used for ranking.

### Pseudotime and trajectories

The pseudotime trajectory analysis was performed using the default pipeline from the *monocle3* package (v1.3.1.)^48^. The original *HuMicA* Seurat object was subset to account for the selected clusters (0, 1, 4, 10, 11). The Seurat object was then converted using the *as.cell_data_set* function from *seurat-wrappers*, and was used as input for trajectory graph learning and pseudotime measurement. The heatmap for the trajectory analysis was developed using *pheatmap*^49^.

### Evaluation of differential cluster composition by pathology

The proportions of each cluster or group of clusters were calculated for each neurologic condition and for each individual subject (Supplementary Figure 4 and Supplementary Table 2). The *sccomp* package^50^ was used to infer the significance of the differential distribution. The *contrasts* parameter was included to estimate depletion or expansion of a cluster in a specific pathology in comparison to the healthy population. The results are presented as the credible interval of the slope and statistical significance is considered for FDR < 0.05.

### Processing of an independent scRNA-seq dataset and external validation

The raw counts matrix and a corresponding cell annotation file were downloaded from https://github.com/vilasmenon/Microglia_Olah_et_al_2020 and the SCT normalization and the aforementioned analysis pipeline was applied. In this case, the *vars.to.regress* function of the SCT normalization was applied to the sample of origin to account for batch effect. The distribution of UMI counts, number of features per cell, percentage of mitochondrial and ribosomal RNA were assessed for quality control. Clustering (res=0.5) outputted 15 clusters from which three were excluded since expression patterns coincided with monocytes, T and B cells and ambiguous cells expressing markers from different CNS cell types. The remaining 12 clusters were used to evaluate the prevalence of the expression patterns described in the *HuMicA*. For GSEA analysis of differential expression of the clusters on the Olah dataset, the results from *FindAllMarkers* (*min.pct = 0.1 and logfc.threshold = 0, test.use=”wilcox”*) were used as input in *fgsea*, and *avg_log2FC* and the p_val_adj were used for ranking. We used the *FindTransferAnchors* function from *Seurat* to perform unsupervised identification of anchors between the Olah dataset (query) and our *HuMicA* (reference). To each *Olah* cell a prediction score was attributed for each of the *HuMicA* subpopulations.

## ACKNOWLEDGEMENTS

We would like to acknowledge all the authors that contributed for the development of the studies from which the data used in our *HuMicA* was collected. The importance of public data access must always be reinforced as a paramount tool for the progression of scientific knowledge, allowing for alternative interpretations of the original data and additional discoveries.

We thank the CERCA Programme/Generalitat de Catalunya, the Josep Carreras Foundation and ICBAS-UP for institutional support. EB is funded by the Spanish Ministry of Science and Innovation (MICINN) [PID2020117212RB-I00; AEI/10.13039/501100011033]. RM-F is funded by an FCT (*Fundação para a Ciência e Tecnologia*) fellowship (SFRH/BD/137900/2018). UMIB is funded by FCT Portugal (UIDB/00215/2020 and UIDP/00215/2020), and ITR (LA/P/006/2020).

Our acknowledgments also extent to all members of the Epigenetics and Immune Disease Group at the Josep Carreras Leukaemia Research institute, to Professor Berta Martins da Silva and the Immunogenetics Laboratory of the Molecular Pathology and Immunology Department of the ICBAS-UP, and to Neurophysiology and Neurology Departments of *Hospital de Santo António – Centro Hospitalar e Universitário do Porto* (HSA-CHUP) represented by Professor António Martins da Silva, MD and Dr. João Chaves for the continuous collaborations and guidance.

## DATA AVAILABILITY

The data and code used in this study is available at https://github.com/RicardoMartins-Ferreira/HuMicA. The integrated *HuMicA* Seurat object is available in an interactive Shiny-based web interface (https://in7yqx-ricardo-ferreira.shinyapps.io/shinyapp/), generated using *ShinyCell*^51^.

## COMPETING INTERESTS

The authors declare no competing interests,

## AUTHOR CONTRIBUTIONS

R.M.-F. and E.B. conceived the study. The bioinformatic analysis was performed by R.M-F. and J.C.-S. performed the bioinformatic analysis. R.M.-F., J.C.-S., J.R.-U., E.M. and E.B contributed for the verification and interpretation of the analysis. R.M.-F. and E.B. were responsible for the writing of the article. All the remaining authors reviewed, edited and approved the final version of the manuscript.

## Notes

### Competing Interest Statement

The authors have declared no competing interest.

## REFERENCES

1. Ginhoux, F. et al. Fate mapping analysis reveals that adult microglia derive from primitive macrophages. Science 330, 841–845 (2010).

2. Ginhoux, F., Lim, S., Hoeffel, G., Low, D. & Huber, T. Origin and differentiation of microglia. Front. Cell. Neurosci. 7, 45 (2013).

3. Martins-Ferreira, R., Leal, B., Costa, P. P. E. & Ballestar, E. Microglial Innate Memory and Epigenetic Reprogramming in Neurological Disorders. Prog. Neurobiol. 101971 (2020) doi:10.1016/j.pneurobio.2020.101971.

4. Subhramanyam, C. S., Wang, C., Hu, Q. & Dheen, S. T. Microglia-mediated neuroinflammation in neurodegenerative diseases. Semin. Cell Dev. Biol. 94, 112–120 (2019).

5. Keren-Shaul, H. et al. A Unique Microglia Type Associated with Restricting Development of Alzheimer’s Disease. Cell 169, 1276–1290.e17 (2017).

6. Habib, N. et al. Massively parallel single-nucleus RNA-seq with DroNc-seq. Nat. Methods 14, 955–958 (2017).

7. Mathys, H. et al. Single-cell transcriptomic analysis of Alzheimer’s disease. Nature 570, 332– 337 (2019).

8. Grubman, A. et al. A single-cell atlas of entorhinal cortex from individuals with Alzheimer’s disease reveals cell-type-specific gene expression regulation. Nat. Neurosci. 22, 2087–2097 (2019).

9. Lau, S.-F., Cao, H., Fu, A. K. Y. & Ip, N. Y. Single-nucleus transcriptome analysis reveals dysregulation of angiogenic endothelial cells and neuroprotective glia in Alzheimer’s disease. Proc. Natl. Acad. Sci. U. S. A. 117, 25800–25809 (2020).

10. Leng, K. et al. Molecular characterization of selectively vulnerable neurons in Alzheimer’s disease. Nat. Neurosci. 24, 276–287 (2021).

11. Morabito, S. et al. Single-nucleus chromatin accessibility and transcriptomic characterization of Alzheimer’s disease. Nat. Genet. 53, 1143–1155 (2021).

12. Zhou, Y. et al. Human and mouse single-nucleus transcriptomics reveal TREM2-dependent and TREM2-independent cellular responses in Alzheimer’s disease. Nat. Med. 26, 131–142 (2020).

13. Jäkel, S. et al. Altered human oligodendrocyte heterogeneity in multiple sclerosis. Nature 566, 543–547 (2019).

14. Schirmer, L. et al. Neuronal vulnerability and multilineage diversity in multiple sclerosis. Nature 573, 75–82 (2019).

15. Velmeshev, D. et al. Single-cell genomics identifies cell type-specific molecular changes in autism. Science 364, 685–689 (2019).

16. Feleke, R. et al. Cross-platform transcriptional profiling identifies common and distinct molecular pathologies in Lewy body diseases. Acta Neuropathol. 142, 449–474 (2021).

17. Masuda, T. et al. Spatial and temporal heterogeneity of mouse and human microglia at single-cell resolution. Nature 566, 388–392 (2019).

18. Wright-Jin, E. C. & Gutmann, D. H. Microglia as Dynamic Cellular Mediators of Brain Function. Trends Mol. Med. 25, 967–979 (2019).

19. Yang, A. C. et al. Dysregulation of brain and choroid plexus cell types in severe COVID-19. Nature 595, 565–571 (2021).

20. Fullard, J. F. et al. Single-nucleus transcriptome analysis of human brain immune response in patients with severe COVID-19. Genome Med. 13, 118 (2021).

21. Lopez-Leon, S. et al. More than 50 long-term effects of COVID-19: a systematic review and meta-analysis. Sci. Rep. 11, 16144 (2021).

22. Huang, L. et al. 1-year outcomes in hospital survivors with COVID-19: a longitudinal cohort study. *Lancet (London*, England) 398, 747–758 (2021).

23. Douaud, G. et al. SARS-CoV-2 is associated with changes in brain structure in UK Biobank. Nature (2022) doi:10.1038/s41586-022-04569-5.

24. Tran, M. N. et al. Single-nucleus transcriptome analysis reveals cell-type-specific molecular signatures across reward circuitry in the human brain. Neuron 109, 3088–3103.e5 (2021).

25. Harris, H. K. et al. Disruption of RFX family transcription factors causes autism, attention-deficit/hyperactivity disorder, intellectual disability, and dysregulated behavior. Genet. Med. Off. J. Am. Coll. Med. Genet. 23, 1028–1040 (2021).

26. Thrupp, N. et al. Single-Nucleus RNA-Seq Is Not Suitable for Detection of Microglial Activation Genes in Humans. Cell Rep. 32, 108189 (2020).

27. Silvin, A. et al. Dual ontogeny of disease-associated microglia and disease inflammatory macrophages in aging and neurodegeneration. Immunity 55, 1448–1465.e6 (2022).

28. Tan, W. et al. Distinct phases of adult microglia proliferation: a Myc-mediated early phase and a Tnfaip3-mediated late phase. Cell Discov. 8, 34 (2022).

29. Ta, H. M. et al. Atf6α deficiency suppresses microglial activation and ameliorates pathology of experimental autoimmune encephalomyelitis. J. Neurochem. 139, 1124–1137 (2016).

30. Olah, M. et al. Single cell RNA sequencing of human microglia uncovers a subset associated with Alzheimer’s disease. Nat. Commun. 11, 6129 (2020).

31. Hammond, T. R. et al. Single-Cell RNA Sequencing of Microglia throughout the Mouse Lifespan and in the Injured Brain Reveals Complex Cell-State Changes. Immunity 50, 253–271.e6 (2019).

32. Li, Q. et al. Developmental Heterogeneity of Microglia and Brain Myeloid Cells Revealed by Deep Single-Cell RNA Sequencing. Neuron 101, 207–223.e10 (2019).

33. Gerrits, E. et al. Distinct amyloid-β and tau-associated microglia profiles in Alzheimer’s disease. Acta Neuropathol. 141, 681–696 (2021).

34. Pappalardo, J. L. et al. Transcriptomic and clonal characterization of T cells in the human central nervous system. Sci. Immunol. 5, (2020).

35. Franjic, D. et al. Transcriptomic taxonomy and neurogenic trajectories of adult human, macaque, and pig hippocampal and entorhinal cells. Neuron 110, 452–469.e14 (2022).

36. Zheng, G. X. Y. et al. Massively parallel digital transcriptional profiling of single cells. Nat. Commun. 8, 14049 (2017).

37. Hao, Y. et al. Integrated analysis of multimodal single-cell data. Cell 184, 3573–3587.e29 (2021).

38. Zhu, Y., Wang, L., Yin, Y. & Yang, E. Systematic analysis of gene expression patterns associated with postmortem interval in human tissues. Sci. Rep. 7, 5435 (2017).

39. McGinnis, C. S., Murrow, L. M. & Gartner, Z. J. DoubletFinder: Doublet Detection in Single-Cell RNA Sequencing Data Using Artificial Nearest Neighbors. Cell Syst. 8, 329–337.e4 (2019).

40. Hafemeister, C. & Satija, R. Normalization and variance stabilization of single-cell RNA-seq data using regularized negative binomial regression. Genome Biol. 20, 296 (2019).

41. Stuart, T. et al. Comprehensive Integration of Single-Cell Data. Cell 177, 1888–1902.e21 (2019).

42. McCarthy, D. J., Campbell, K. R., Lun, A. T. L. & Wills, Q. F. Scater: pre-processing, quality control, normalization and visualization of single-cell RNA-seq data in R. Bioinformatics 33, 1179–1186 (2017).

43. Alquicira-Hernandez, J. & Powell, J. E. Nebulosa recovers single cell gene expression signals by kernel density estimation. Bioinformatics (2021) doi:10.1093/bioinformatics/btab003.

44. Wu, T. et al. clusterProfiler 4.0: A universal enrichment tool for interpreting omics data. Innov. (Cambridge 2, 100141 (2021).

45. Garcia-Alonso, L. et al. Transcription Factor Activities Enhance Markers of Drug Sensitivity in Cancer. Cancer Res. 78, 769–780 (2018).

46. Alvarez, M. J. et al. Functional characterization of somatic mutations in cancer using network-based inference of protein activity. Nat. Genet. 48, 838–847 (2016).

47. Korotkevich, G. et al. Fast gene set enrichment analysis. bioRxiv 60012 (2021) doi:10.1101/060012.

48. Cao, J. et al. The single-cell transcriptional landscape of mammalian organogenesis. Nature 566, 496–502 (2019).

49. Kolde, R. pheatmap: Pretty Heatmaps. R package version 1.0.12 https://cran.r-project.org/package=pheatmap (2019).

50. Mangiola, S. sccomp: Robust Outlier-aware Estimation of Composition and Heterogeneity for Single-cell Data. R package version 1.2.1 https://github.com/stemangiola/sccomp (2022).

51. Ouyang, J. F., Kamaraj, U. S., Cao, E. Y. & Rackham, O. J. L. ShinyCell: simple and sharable visualization of single-cell gene expression data. Bioinformatics 37, 3374–3376 (2021).

