## Supplementary Figure Legends for "The Human Microglia Atlas (HuMicA) Unravels Changes in Homeostatic and Disease-Associated Microglia Subsets across Neurodegenerative Conditions"

**Supplementary Figure 1.** UMAP visualization of the six main cell types annotated for the Mathys **(A)**, Grubman **(B)**, Leng **(C)**, Morabito **(D)**, Lau **(E)**, Zhou **(F)**, Pappalardo **(G)**, Thrupp **(H)**, Jakel **(I)**, Schirmer **(J)**, Velmeshev **(K)**, Feleke **(L)**, Tran **(M)**, Franjic **(N)**, Yang **(O)** and Fullard **(P)** datasets.

**Supplementary Figure 2. (A)** Expression of canonical cell type markers in the subset of immune cells extracted from each dataset (*PLP1*, oligodendrocytes; *SNAP25*, neurons; *DOCK8*, immune cells; *VCAN*, OPCs; *SLC1A2*, astrocytes; *FLT1*, endothelial cells). **(B)** UMAPs showing the module score (MSc) expression of several immune cell/microglia markers (*P2RY12, DOCK8, CD74, CX3CR1, C1QB, C3, AIF1, HLA-DRA*) in each dataset. Expression is prevalent in the clusters annotated as immune cells.

**Supplementary Figure 3.** Evaluation of covariate confounding effects. Nuclei were annotated in the UMAP plots in relation to *Study* **(A)**, *Tissue region* **(B)**, *Age* **(C)**, *Group* **(D)**, *Gender* **(E)** and *Tissue state* **(F)**. For each category, the number of nuclei and the mean genes detected per nuclei are presented in a barplot. **(G)** Density plot representation of the percentage of variance explained by each of the covariates.

**Supplementary Figure 4.** Proportion distribution of all clusters in the *HuMicA* across pathological conditions as a whole **(A)** and individualized by subject **(B).**

**Supplementary Figure 5. (A)** UMAP representation of the *HuMicA* clusters 3, 6 and 7. **(B)** Dot plot representation of the expression of markers of neurons (*SYT1, SNAP25*) and oligodendrocytes (*PLP1* and *ST18*)*,* and module score (MSc) expression of the lists of genes upregulated in clusters 3, 6 and 7 vs all other microglia. **(C)** Differential distributions of clusters 3, 6 and 7 in each pathology in relation to the healthy group. Population expansion or depletion is represented by the credible interval of the slop. Statistical significance is considered for an adjusted p value (FDR) < 0.05 and is highlighted in red. **(D)** Overlap plots of the enriched gene ontology (GO) terms calculated for the upregulated DEGs between all microglia clusters. Highlighted by the dashed red boxes are the overlapping GO terms between cluster 10 (*Homeos3*) and the other homeostatic clusters.

**Supplementary Figure 6. (A)** UMAPs showing the expression of genes specifically upregulated in each homeostatic cluster (*NAV2*, *Homeos1*; *GRID2*, *Homeos2*; *SERPINE1*, *Homeos3;* *CCDC26*, *Homeos4*). **(B)** Representation of the top ten most significantly enriched GO terms considering the upregulated DEGs in homeostatic clusters vs other microglia and specifically in each of the homeostatic subpopulations. **(C)** TF enrichment of all significant TFs (p value < 0.01) to the comparison of differential expression of all homeostatic clusters vs all other microglia, and specifically in each of the homeostatic subpopulations. Significance is represented by the Normalized Enrichment Score (NES) and the negative of log of the adjusted p value (FDR). **(D)** Overlap plots of the upregulated and downregulated DEGs calculated for the homeostatic microglia in each pathology vs the healthy population. The bars highlighted in gold (upregulated) and purple (downregulated) represent the DEGs exclusive to each pathology. **(E)** Proportions of all homeostatic microglia across pathological conditions as a whole and individualized by subject. Dashed line represents the proportion in the healthy group (“*No Neuropathology”*). **(F)** Proportions of each homeostatic cluster across pathological conditions as a whole and individualized by subject. Dashed line represents the proportion in the healthy group (“*No Neuropathology”*).

**Supplementary Figure 7. (A)** UMAPs showing the gene expression of the module score (MSc) of the conserved DIM human signature (*CCL4, CD14, CD83, CSF2RA, EIF1, FOS, IER2, JUN, JUNB, EGR1, IL1B, TNF, PLAUR, SAT1, ZFP36, BTG2*) from Silvin *et al.* (2022)^27^, and the individual expression of *CD83, CSF2RA* and *SAT1.* **(B)** Net plot representation of the fifteen most significant GO terms enriched on the lists DEGs in the *DIM* vs *DAM* comparison. The range of expression of each gene is represented by the negative of the log of the p value and the average of the log2 of the fold change. High expression (red) corresponds to genes upregulated in *DIM* and low expression (blue) corresponds to genes upregulated in *DAMs* **(C)** Barplot representing all the significantly enriched GSEA terms from the *REACTOME, WILKIPATHWAYS* and *KEGG* repositories for the differential expression of *DIM* vs *DAM* populations. Enrichment is represented as a function of the negative of log of the adjusted p value and Normalized Enrichment Score (NES).

**Supplementary Figure 8. (A)** UMAPs showing the module score (MSc) expression of the set of genes characteristic of the stage 1 DAM phenotype (*APOE, B2M, TYROBP, FTH1, LYZ, CTSB, CTSD*), and the individual expression of *APOE, B2M* and *FTH1*; and the MSc expression of the set of genes characteristic of stage 2 DAM phenotype (*LPL, CTSL, TREM2, AXL, CD9, CSF1, CCL23, ITGAX, CLEC7A, LILRB4, TIMP2*), and the individual expression of *SPP1, ITGAX* and *SLC11A1*. **(B)** Net plot representation of the fifteen most significant GO terms enriched on the lists of DEGs in the *Final.DAM* vs *Intermediate.DAM* comparison. The range of expression of each gene is represented by the negative of the log of the p value and the average of the log of the fold change. High expression (red) corresponds to genes upregulated in *Final.DAM* and low expression (blue) corresponds to genes upregulated in *Intermediate.DAM*. **(C)** Barplot representing all the significantly enriched GSEA terms from the *REACTOME, WILKIPATHWAYS* and *KEGG* repositories for the differential expression of *Final.DAM* vs *Intermediate.DAM* populations. Enrichment is represented as a function of the negative of log of the adjusted p value and Normalized Enrichment Score (NES).

**Supplementary Figure 9. (A)** Representation of the enrichment of all significant TFs (p value < 0.01) relative to the comparison of differential expression of *DAM* vs other microglia, *Final.DAM* vs *Intermediate.DAM*, *DIM* vs other microglia and *DIM* vs *DAM*. Significance is represented by the NES (Normalized Enrichment Score) and the negative of log of the adjusted p value (FDR). **(B)** Heatmap representation of the z-score of the average expression (by pathological condition) of the top ten most upregulated genes in all *DAM* nuclei and in *DIM* that are exclusive to each pathology when compared with the healthy group; coupled with the overlap plots for the upregulated and downregulated DEGs calculated for each pathology vs the healthy population. The bars highlighted in gold (upregulated) and purple (downregulated) represent the DEGs exclusive to each pathology. **(C)** Differential distribution of each DAM and DIM clusters in each pathology in relation to the healthy group. Population expansion or depletion is represented by the credible interval of the slop. Statistical significance is considered for an adjusted p value (FDR) < 0.05 and is highlighted in red.**(D)** Proportions of all *DAM* nuclei and of the individual *Final.DAM*, *Intermediate.DAM* and *DIM* subpopulations across pathological conditions as a whole and individualized by subject. Dashed line represents the proportion in the healthy group (“*No Neuropathology*”).

**Supplementary Figure 10. (A)** Dot plot representation of the top five most upregulated genes (ordered by avg_log2FC) for each of the subpopulations within the *Final.DAM*, *Intermediate.DAM* and *DIM* clusters. **(B)** Proportions of all subpopulations across pathological conditions as a whole. Dashed line represents the proportion in the healthy group (“*No Neuropathology*”). **(C)** GO enrichment of the top three (ordered by the negative of logFDR and logFC) obtained from the lists of upregulated DEGs in each subpopulation.
